# A multi-omics analysis of *Arabidopsis thaliana* root tips under Cd exposure: A role of HY5 in limiting accumulation

**DOI:** 10.1101/2024.08.29.609871

**Authors:** Ludwig Richtmann, Noémie Thiébaut, Alok Ranjan, Manon Sarthou, Stéphanie Boutet, Marc Hanikenne, Stephan Clemens, Nathalie Verbruggen

## Abstract

- Cadmium (Cd) is a major environmental pollutant with high toxicity potential. Even though a reduction of growth, including the primary root, is a clear consequence of Cd exposure, a profound understanding of the impact of Cd on the root apical meristem (RAM) and the elongation/differentiation zone (EDZ) is still lacking.
- In this study, *Arabidopsis thaliana* roots were subjected to Cd and divided into root tips (RT) and remaining roots (RR) to separately assess the effect of Cd using transcriptomics, ionomics and metabolomics.
- Elemental profiling revealed lower Cd accumulation in RT and differences in mineral contents between RT and RR. Transcriptomic analysis demonstrated distinct gene expression patterns in RT and RR, with Cd having less impact in RT. Functional enrichment analysis revealed genes associated with iron and sulfur homeostasis as well as the response to light in both RR and RT. RT-specific responses to Cd included several genes regulated by the transcription factor ELONGATED HYPOCOTYL 5 (HY5) and notably, the *hy5* mutant showed increased Cd sensitivity and accumulation compared to the wild type.
- This study provides comprehensive insights into the inhibitory effects of Cd on primary root growth, elucidating molecular mechanisms involved, particularly highlighting the role of HY5 in Cd accumulation.

## Introduction

In natural environments, plant roots encounter diverse edaphic conditions and are confronted with variations in soil texture, water saturation, pH levels, salinity or nutrient availability. A variety of metal ions are required as macro-(Ca, K, Mg) or micronutrients (Fe, Cu, Co, Mn, Mo, Ni, Zn) and their availabilities in habitats nearly always deviate from optimal conditions (Lin & Aarts, 2012; Kumar *et al*., 2021). Therefore, plants need a tightly regulated metal homeostasis network in order to handle either deficiencies or excess of metals, with the latter potentially causing toxicity. Apart from that, metals such as cadmium (Cd) are present in soil as environmental contaminants. They are not essential for plant growth and can be highly toxic (Lin & Aarts, 2012; Haider *et al*., 2021).

As a metal with known carcinogenic potential, the presence of Cd in agricultural soils, together with its uptake by crops additionally represents a substantial threat to human health and motivates studies on the interaction of Cd with plants (Clemens *et al*., 2013; Rahim *et al*., 2022).

Plant root responses to maintain elemental homeostasis in challenging soil conditions can be broadly categorized into two general types. On the one hand, acclimatization to such environmental conditions can occur through developmental plasticity. For instance, *Arabidopsis thaliana* seedlings exhibit notable morphological alterations when grown under varying phosphate (P_i_) availabilities. These involve changes in root architecture such as the inhibition of primary root growth and enhanced elongation of the lateral roots and root hairs (Ren *et al*., 2023). On the other hand, plants acclimatize to soil environments by modulating molecular networks that regulate uptake, transport, and storage of metals and other nutrients. As a response to iron (Fe) deficiency for example, the transcriptional upregulation of central Fe uptake genes such as *IRT1* and *FRO2* relies on the formation of heterodimeric complexes between FIT and the group Ib basic helix-loop-helix (bHLH) transcription factors bHLH38 and bHLH39 (Riaz & Guerinot, 2021).

In plants, the most prominent developmental consequence of Cd exposure is a reduction of growth in both above- and below-ground organs. This is the result of diverse toxicity mechanisms including the inhibition of enzymes, lipid peroxidation, overproduction of reactive oxygen species (ROS), induction of DNA damage and reduction of photosynthetic activity (Lin & Aarts, 2012; Haider *et al*., 2021). To mitigate toxic effects upon Cd entry into the plant, the presence of efficient detoxification mechanisms is crucial. This can be exemplified by the extreme Cd-hypersensitivity of the *cad1-3* mutant. The *AtPCS1* allele of *cad1-3* strongly impairs its ability to synthesize metal-chelating phytochelatins (PCs), which, together with vacuolar sequestration of PC-Cd complexes, is a key mechanism to cope with metals such as Cd (Howden *et al*., 1995; Lin & Aarts, 2012).

Besides these toxicity mechanisms, Cd is known to interfere with the homeostasis of essential elements. One example is the intricate relationship with Fe acquisition (Wu *et al*., 2012; Zhai *et al*., 2014; Zhou *et al*., 2021; Wang *et al*., 2023). Cd exposure was shown to result in a reduction of root Fe content and the activation of Fe-deficiency responses (Lešková *et al*., 2017). At least in part, the connection between Cd and Fe homeostasis may be attributable to shared uptake pathways of the two metals. For instance, multiple studies suggest that, in *A. thaliana*, IRT1, a member of the ZIP (ZRT- and IRT-like proteins) family, plays a significant role in Cd uptake (Connolly *et al*., 2002; Vert *et al*., 2002; Fan *et al*., 2014; Ismael *et al*., 2019). Additionally, in the hypertolerant species *Arabidopsis halleri*, ZIP6 was shown to be a Cd-uptake transporter (Spielmann *et al*., 2020). Besides ZIP family proteins, a significant contribution to Cd uptake under low nitrate conditions was demonstrated for the nitrate transporter NRT2.1 (Guan *et al*., 2021). Furthermore, there is evidence for a major role of the manganese (Mn) and Fe transporter NRAMP5 in Cd uptake in rice (Ishimaru *et al*., 2012; Clemens *et al*., 2013).

Due to the abovementioned toxicity mechanisms, elevated Cd concentrations in the rhizosphere affect many aspects of root anatomy such as lateral root emergence (Wang *et al*., 2021), root thickness (Lux *et al*., 2011), root hair density (Bahmani *et al*., 2022) and primary root growth. The latter depends on the tissues in the root apex: the root apical meristem (RAM), where cells undergo mitotic divisions, and the elongation/differentiation zone (EDZ), where they expand to their final size and differentiate to achieve tissue-specific characteristics. The two zones are separated by the transition zone (TZ). Optimal root growth depends on meristem maintenance, which represents a tight balance between division and elongation/differentiation rates (Perilli *et al*., 2012).

This balance is regulated, among others, by the two phytohormones auxin and cytokinin and the cellular response to their distribution in the root plays a critical role for the developmental characteristics in the different zones of the root apex (Dello Ioio *et al*., 2007; Růžička *et al*., 2009; Perilli *et al*., 2010; Jia *et al*., 2023; Tao *et al*., 2024). Independently of auxin and cytokinin hormonal signaling, the developmental processes in the root tip are controlled by the distribution of reactive oxygen species (ROS). Proliferation and differentiation rates depend on a gradient between superoxide ions (O_2_^•−^) and hydrogen peroxide (H_2_O_2_) (Tsukagoshi *et al*., 2010; Wells *et al*., 2010).

Several studies have addressed the impact of Cd on the root apex of *A. thaliana*, examining disruptions in the auxin/cytokinin hormonal balance (Yuan & Huang, 2016; Bruno *et al*., 2017; Leonardo *et al*., 2021) or the effects on the cell cycle (Cui *et al*., 2017). However, the effects of metal stress on the molecular networks regulating root growth and nutrient homeostasis have not been comprehensively examined.

Therefore, this study followed a multidirectional omics approach in *A. thaliana* plants exposed to Cd to investigate i) the transcriptomic response, ii) metal homeostasis and iii) the metabolic landscape in the root tips (RT) compared to remaining roots (RR).

Our analysis revealed lower Cd accumulation in RT compared to RR and highlighted differences between the two root parts. We additionally provided insight into general differences in gene expression between RT and RR. Notably, the transcriptomic data indicated a specific involvement of the transcription factor ELONGATED HYPOCOTYL 5 (HY5) in the response of RT to Cd. Phenotypic analysis of two loss of function *hy5* mutants confirmed the functional relevance of HY5 in the response to Cd, as these mutants displayed increased Cd sensitivity which was accompanied by elevated cellular Cd levels compared to wild type plants.

## Materials & Methods

### Plant material & growth conditions

*A. thaliana* (Col-0) seeds were surface sterilized by rinsing 3 min with 70% ethanol and 2 min with 2% bleach. Upon 3 washing steps with sterile H_2_O, the seeds were suspended in 0.1% (w/v) sterile agar. All seeds were sawn on nylon meshes and stratified (4°C, 2 days).

Plants were grown vertically on agar plates with Hoagland medium (1.5mM Ca(NO_3_)_2_, 0.28mM KH_2_PO_4_, 0.75mM MgSO_4_, 1.25mM KNO_3_, 0.5μM CuSO_4_, 1μM ZnSO_4_, 5μM MnSO_4_, 25μM H_3_BO_3_, 0.1μM Na_2_MoO_4_, 50μM KCl, 10μM Fe-HBED, 0.5g*l^-1^ MES, 1% (w/v) sucrose, 0.8% (w/v) agar (M-type, Sigma), pH 5.7) in a CLF Plant Climatics GroBank under long day light conditions (16h light, 21°C, 100PAR/ 8h dark, 18°C). Fe-deficiency assays were conducted by adding 75µM FerroZine. The mutant lines used were SALK_066689C (*xth20)*, SALK_042683C (*xth26)*, SALK_001585 (*cyp82c4),* GK_578C04 *(hrg1),* GK_772A03 *(hrg2)*, GABI DUPLO 1202/1a1.16.1 *(elip1/2,* NASC ID N2103098), *myb12-1-f* (Mehrtens *et al*., 2005), SALK_0966551C (herein called *hy5_1)* and *hy5_215* (Oyama *et al*., 1997).

### ICP-MS

24h and 48h after transferring 7-day-old seedlings to Cd-contaminated medium (25µM with CdCl_2_), plants were desorbed prior to ICP analysis. Therefore, they were washed twice in a desorption solution [(5mM CaCl_2,_ 1mM MES, pH5.7), 10min, 4°C] and twice in distilled water (5min, 4°C). RT samples were collected by separating ∼2-3mm root tips (RT) from the remaining root (RR). Dry root material was weighed into polytetrafluoroethylene (PTFE) digestion tubes and concentrated nitric acid (0.5ml, 67-69%) was added to each tube. After 4h incubation, samples were digested under pressure using a high-performance microwave reactor (Ultraclave 4, MLS). Digested samples were transferred to Greiner centrifuge tubes and diluted with de-ionized water. Elemental analysis was performed using inductively coupled plasma - mass spectrometry (Sector Field High Resolution (HR)-ICP-MS, ELEMENT 2, Thermo Fisher Scientific). For sample introduction a SC-2 DX Autosampler (ESI, Elemental Scientific) was used. A 6-point external calibration curve was set from a certified multiple standards solution (Bernd Kraft). The elements rhodium (Rh) and germanium (Ge) were infused online and used as internal standards.

### RNA isolation, cDNA synthesis & RNA-Sequencing

Seedlings were grown for 7d on control medium before being transferred to Cd contaminated medium (25µM). 24h and 48h after transferring seedlings, samples were collected by cutting RT ∼2-3 mm from the apex, as presented in the companion paper (Thiébaut *et al*.). From each plate, the RR after cutting were also collected. RNA isolation was done with a Maxwell 16 device (Promega) using the Maxwell RSC Plant RNA Kit. This method isolates total RNA using cellulose-based paramagnetic particles and includes a DNAse I-treatment.

RNA quality and concentration were measured with an Agilent 2100 Bioanalyzer Expert using the Agilent Nano 6000 Kit. Library preparation was done with the TruSeq Stranded mRNA Sample Preparation Kit, using 1μg of RNA as starting material. Library quality was assessed with a QIAxcel screening kit. Quantification, dilution and pooling of the libraries was done using the KAPA SYBR® FAST Universal qPCR Kit and an Applied Biosystems 7900HT Real-Time PCR system. Sequencing (2x150bp paired-end reads) was done on a NovaSeq6000 with standard workflow.

### RNA-Seq data processing

Data quality assessment & trimming was done with FastQC & Trimmomatic (Bolger *et al*., 2014). HISAT2 was used for mapping (TAIR10 genome assembly and the Araport11 annotation were used (Cheng *et al*., 2017)). Read counting was done with htseq-count. Differential expression analysis was performed with DESeq2 (Love *et al*., 2014). DEG were extracted with log2_FoldChange_ > 1 or log2_FoldChange_ < -1 and p_ajd_ < 0.05. For GO overrepresentation analysis, the online tool from the PANTHER website (pantherdb.org, June 2022) was used.

### Metabolomic analysis

For untargeted metabolomic analysis, plants were grown as described above. 48h after transferring seedlings, RT and RR samples were taken by cutting and collecting similar sized pieces (∼2-3mm) of RT and RR (at ∼mid-length of the primary root). Extraction and LC-MS/MS analysis of polar/semipolar primary and specialized metabolites from root material was done as described in Boutet *et al*., (2022). For comparing the quantity of metabolites between RR and RT, metabolite concentrations were normalized to the amount of collected root pieces for each sample.

For further analysis, relative metabolite quantifications were normalized to the total metabolite count of each sample. Statistical analysis was done by fitting a linear model to log2 transformed metabolite data with limma and applying empirical Bayes moderation of the linear model fit. Features were considered as significant DAF (differentially abundant features) when log2_FoldChange_ > 1 or log2_FoldChange_ < -1 and p_ajd_ ≤ 0.05.

### ICP-OES

Roots and shoots of seedlings were desorbed rinsed with Milli-Q water (10min, 4°C), washed twice with 20mM CaCl_2_ (10min, 4°C), once with 10mM ethylenediaminetetraacetic (EDTA) (10min, 4°C, pH5.7) and rinsed one more time with Millipore water (10min, 4°C). Samples were dried, weighed into PTFE reaction tubes and digested in a microwave oven (START 1500, MLS, Leutkirchen) using a 2:1 mixture of HNO_3_ (65%, v/v) and H_2_O_2_ (30%, v/v). Element concentrations were measured with an iCAP 6500 Series ICP-OES (Thermo-Fisher Scientific).

### Thiol profiling with HPLC

For HPLC analysis of thiols, seedlings were grown for 9d on control plates, transferred to control/Cd plates and harvested after 3d. Measurements of phytochelatins with HPLC and glutathione were done as described in Pischke *et al*., (2022).

## Results

### The effect of Cd on primary root growth

To assess the effect of Cd on primary root growth, *A. thaliana* seedlings were grown for 7d on control medium and then transferred to either Cd-containing medium or control medium. Primary root elongation was significantly reduced after 24h and 48h when the plants were exposed to 25µM Cd. After one week under these conditions, root growth was inhibited by 50 % (**Figure 1, A**).

**Figure 1:**
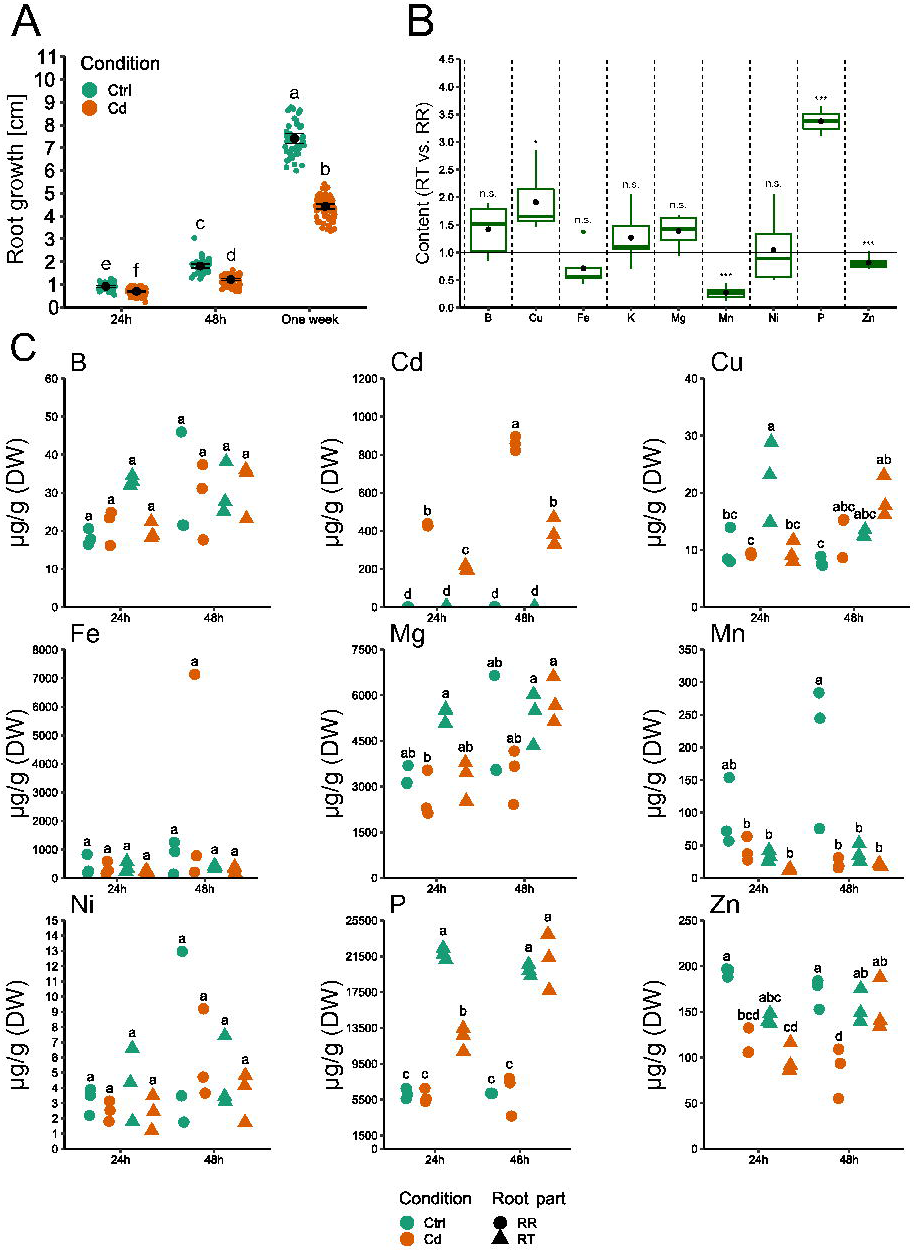
Impact of 25µM Cd on root growth and elemental concentrations in *A. thaliana*. **A.** Primary root growth inhibition by Cd. Seven-day-old seedlings were transferred to control/25µM Cd plates. The progression of growth was monitored after 24h, 48h and one week. Letters indicate statistical significance (ANOVA with Tukey HSD, p ≤ 0.05, n=46-76, data from three independent biological replicates, values are mean +/- SD) **B.** Comparison of elemental concentrations in root tips (RT) versus remaining roots (RR) in control conditions. Asterisks indicate statistical significance (Mann-Whitney U-test, p ≤ 0.05, data from control 24h and control 48h were pooled and averaged, n=6). **C.** Elemental concentrations in root tips (RT) and remaining roots (RR) after Cd exposure. Letters indicate statistical significance (ANOVA with Tukey HSD, p ≤ 0.05, n=3, data from three independent biological replicates).

### Elemental profiles of root tips and remaining roots under control condition and after cadmium exposure

We assessed differences in the mineral concentrations between the root tips (RT) and the remaining roots (RR) in control conditions. Cu and P concentrations were significantly higher in RT compared to RR. In contrast, Mn and Zn concentrations were lower in RT compared to RR (**Figure 1, B**).

As Cd exposure is known to interfere with plant nutrient homeostasis (Haider *et al*., 2021), we analyzed Cd accumulation and its impact on mineral profiles of RT and RR at both 24h and 48h post Cd exposure. We observed an increase in Cd accumulation from 24h to 48h in both RR and RT. However, the Cd concentration in RT was consistently lower than in RR at both timepoints, with RT accumulating only 45-47% the amount of Cd found in RR (**Figure 1, C**). Mn concentrations were reduced after 48h of Cd treatment in RR. Additionally, the concentrations of Cu and P decreased after 24h in RT. Zn concentrations decreased in all Cd-treated samples compared to controls, except in RT at 48h. Cd is known to interfere with Fe homeostasis (Lešková *et al*., 2017). However, due to large variations, no significant difference in Fe concentrations between Cd-treated and control samples were detected in the ICP-MS measurements (**Figure 1, C**). We repeated the measurements on roots of Cd treated seedlings with ICP-OES and found a reduction in Fe concentrations upon Cd exposure in Col-0 (**Supplementary Figure S1**).

### Transcriptomic analysis of *A. thaliana* roots under Cd-stress

We performed a transcriptomic analysis of RT and RR after 24h and 48h of treatment, coinciding with the observations of the effect of Cd on root growth and mineral homeostasis.

By analyzing the influence of the growth conditions, harvesting timepoints and the sampled root parts on global gene expression profiles with principal component analysis (PCA), we identified the root part as the primary factor contributing to transcriptional variance. RT and RR samples were clearly separated along PC1, which explained 96% of the total variance observed. By contrast, the condition (Control vs. Cd) separated samples along PC2, explaining 2% of variance (**Figure 2, A**).

**Figure 2:**
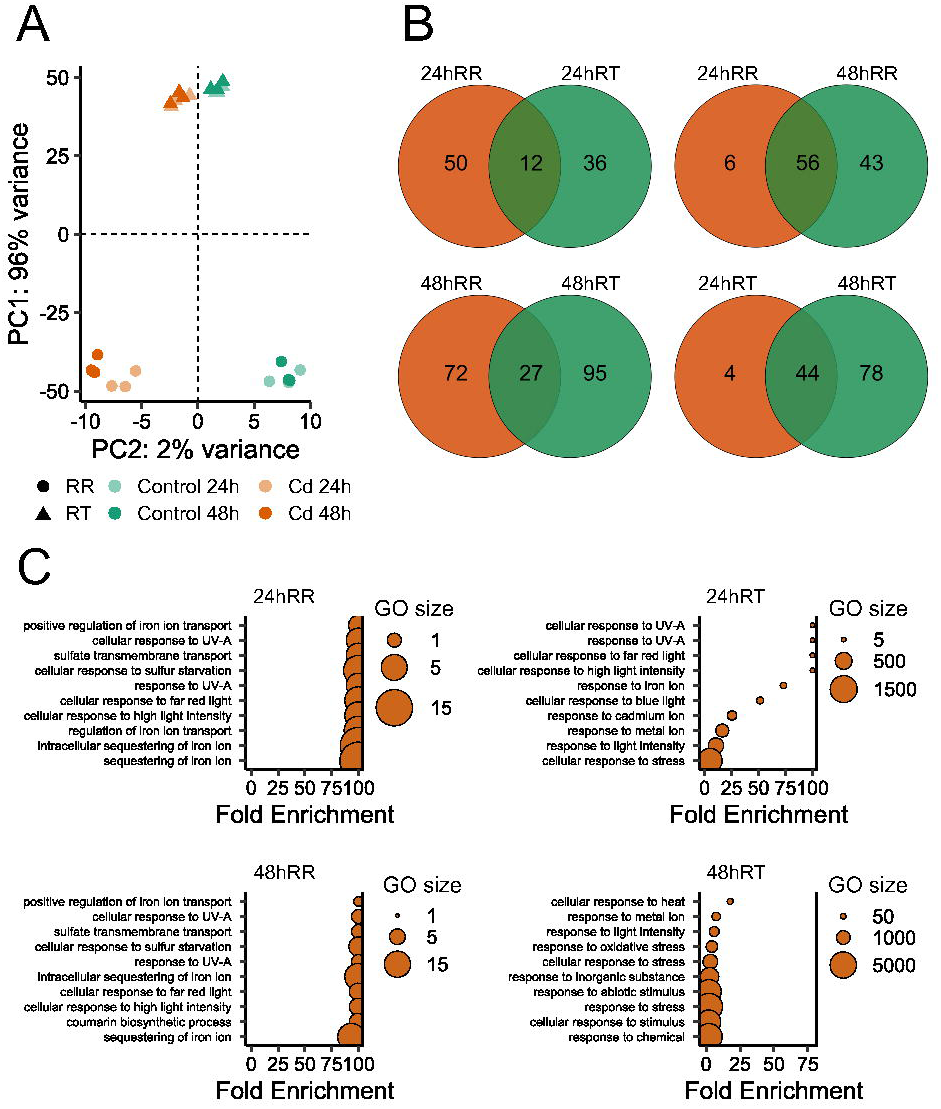
RNA sequencing of *A. thaliana* root tips (RT) and remaining roots (RR) after exposure to 25µM Cd. Seven-day-old seedlings were transferred to control/25µM Cd plates and exposed for 24h or 48h. **A.** Principal component analysis showing the distribution of samples according to PC1 and PC2. The percentage of variance explained by each PC is indicated. **B.** Venn diagrams showing the number of DEG (Control vs. Cd) and the overlap between the different sample types. **C.** GO overrepresentation analysis with the DEG for each timepoint and root part. The amount of genes concerned for each GO term (GO size) and respective fold enrichment are indicated. GO analysis was done with PANTHER. The 10 GO terms with highest enrichment were selected for the graphs.

Since large differences between transcriptomes of RR and RT were identified by PCA, variation of gene expression in the two root parts was explored with greater depth. Analysis of pooled control samples (24h+48h) revealed that 1326 genes and 4544 genes showed higher or lower expression respectively, in RT compared to RR (**Figure 3, B; Supplementary Tables S5 and S6**). Among genes more expressed in RT than in RR, many were related to cellular processes such as ribosome assembly and protein processing/modification. Genes encoding cell-wall modifying proteins with xyloglucan/xyloglucosyl transferase activity were also overrepresented (**Figure 3, A**).

**Figure 3:**
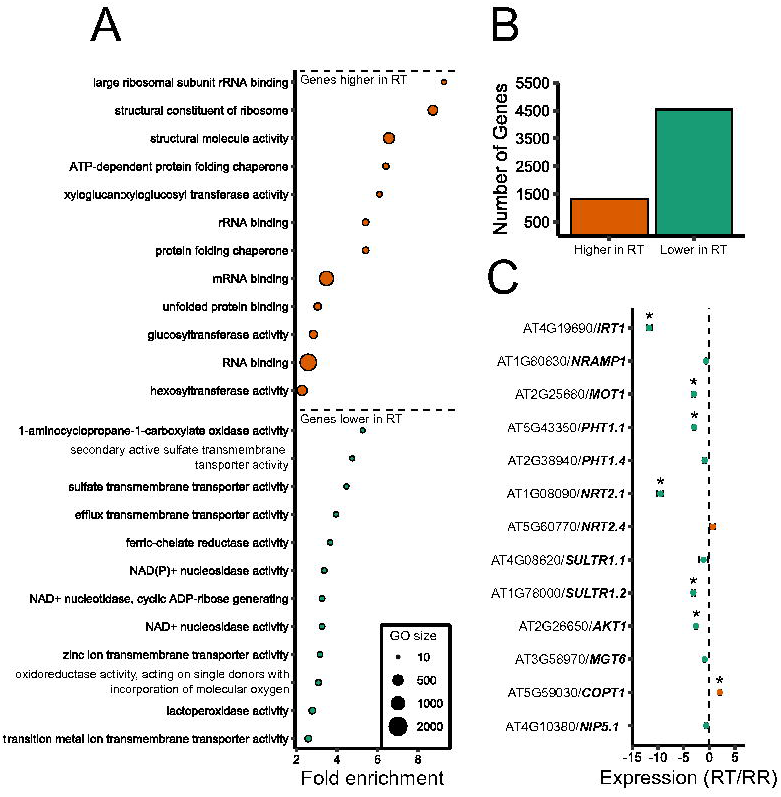
Gene expression in RT vs RR. DEG were extracted from control samples (24 + 48h pooled). **A.** GO overrepresentation analysis made with PANTHER. The size of the GO terms is also shown. **B.** Number of DEGs that are higher or lower in RT compared to RR. **C.** Expression of a selection of genes acting in nutrient uptake in RT vs. RR. Displayed are mean values of log2 transformed fold changes ± standard deviation, 6 independent replications).

On the contrary, genes implicated in ethylene synthesis were less expressed in RT than in RR. Additionally, a large number of differentially expressed genes (DEGs) between RT and RR were related to transmembrane transport activity. Together with the finding of lower Cd accumulation in RT, this prompted us to investigate the expression of metal and nutrient transporters. Many were expressed to a lower extent in RT, including the Fe uptake transporter *IRT1* and the nitrate transporter *NRT2.1*, which are both involved in Cd uptake (Vert *et al*., 2002; Guan *et al*., 2021). Conversely, the copper transporter *COPT1* was found to be more expressed in RT compared to RR (**Figure 3, C**).

Notably, the Cd treatment had a lower impact on gene expression in RT, compared to the RR samples (**Figure 2, A**). Differential expression analysis yielded 50 DEGs 24h after Cd stress in RR and 36 in RT, with an overlap of 12 genes (**Figure 2, B, left**). After 48h of Cd stress, 72 genes were differentially expressed in RR and 95 in RT, with an overlap of 27 genes in the two groups. In RR, 56 genes were differentially expressed at both timepoints, 6 genes exclusively after 24h and 43 genes after 48h. In RT, a similar trend was observed, with an overlap of 44 DEGs at both timepoints and 4 and 78 DEGs exclusively at time points 24h and 48h, respectively (**Figure 2, B, right**).

As one of our goals was to analyze whether elements of the cell cycle regulatory machinery in the root meristem are Cd responsive, we examined the expression of a set of cell-cycle related genes (Vandepoele *et al*., 2002) in RT. However, none of these genes were differentially expressed after Cd-exposure (**Supplementary Figure S2**).

In order to gain insight into the functional categories represented among DEGs in RR and RT, gene ontology (GO) overrepresentation analysis was performed.

The analysis of the response to Cd in RR at both timepoints revealed an enrichment of processes related to Fe homeostasis and sulfur assimilation (**Figure 2, C, left panels**). Transcriptomic changes related to Fe homeostasis included an upregulation of *BASIC HELIX-LOOP-HELIX 38, 39 and 100, IRONMAN 1, 2, 5 and 7, FERRIC REDUCTION OXIDASE 3 and 5* and a downregulation of *VACUOLAR IRON TRANSPORTER-LIKE 1, 2 and 5* and *FERRITIN 1 and 4*. Interestingly *S8H* and *CYP82C4*, which are involved in the coumarin biosynthetic pathway and thus in Fe acquisition (Robe *et al*., 2021a), were found to be downregulated after Cd exposure. Enrichment of DEGs involved in sulfur homeostasis were also identified in Cd-treated RR but not in RT, including *RESPONSE TO LOW SULFUR 1, 2* and *3* as well as *SULPHATE TRANSPORTER 1;1*. In both RR and RT, enriched GO terms related to the response to light also included the genes *EARLY LIGHT INDUCIBLE 1 and 2*, *ELIP1* and *ELIP2* **(Supplementary Tables S1 – S4)**.

The functional categories of DEGs in RT appeared to be related to broader biological functions, but the GO terms were characterized by lower fold enrichment than in RR. Several GO terms related to the response to irradiations as well as to the response to Fe, Cd and metal ions in general were identified (**Figure 2, C, right panels)**.

### The response of root tips to Cd

Exploration of the RT-specific Cd response unveiled several *XTH* (*XYLOGLUCAN ENDOTRANSGLUCOSYLASE/HYDROLASE*) family genes such as *XTH12, 13; MYB12;* and the two *HYDROGEN PEROXIDE RESPONSIVE GENES, HRG1* and *2,* (**Figure 4, A; Supplementary Tables S3 and S4**). Additionally, the *ELIP1* and *ELIP2* were significantly upregulated in both RR and RT, with a notably higher induction in RT upon Cd exposure (**Figure 4, A**).

**Figure 4:**
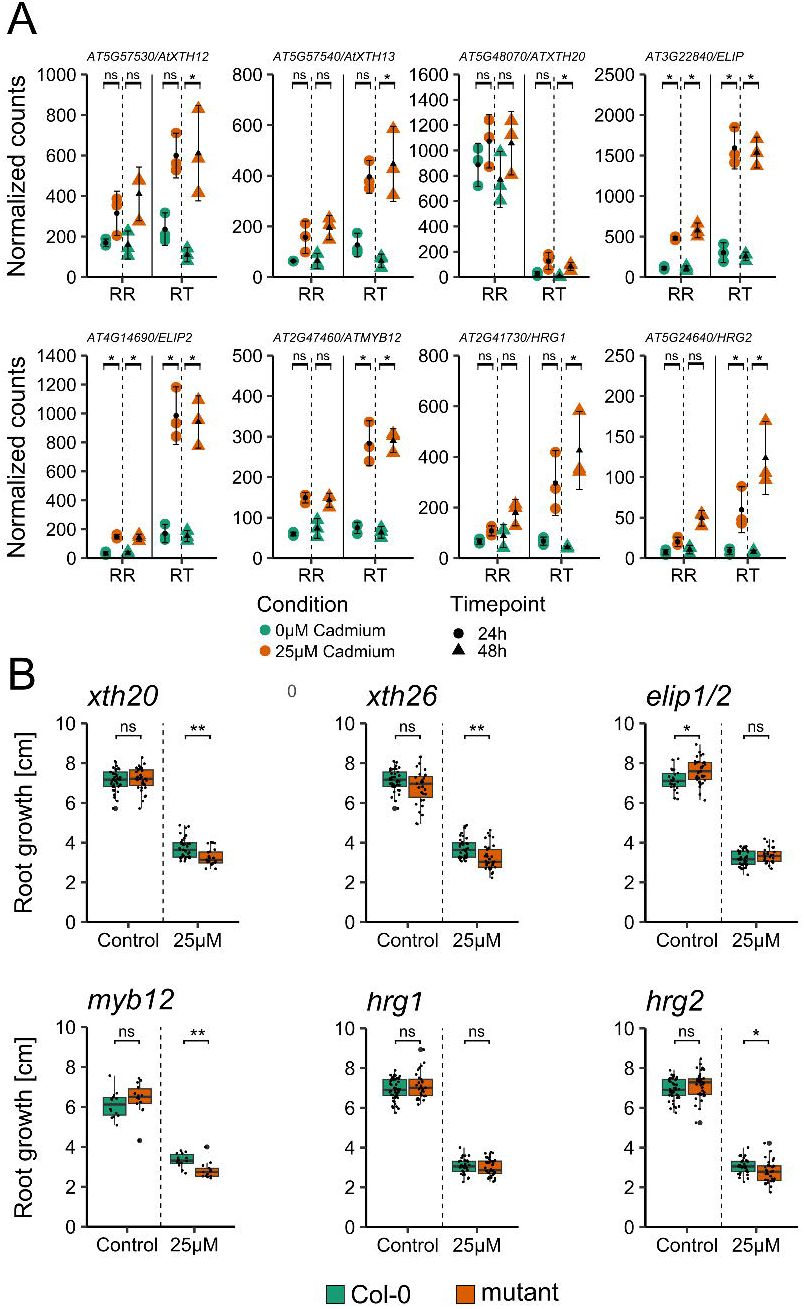
Gene Expression and phenotypic analysis of root tip responsive genes in *A. thaliana*. **A.** Expression of genes that display a specific or more pronounced response to Cd in RT compared to RR. Displayed are mean values ± standard deviation. Asterisks indicate statistical significance of DEG (DESeq2, padj ≤0.05, lfcThreshold=1, data from three independent replications). **B.** Primary root growth of mutants corresponding to the genes displayed in **A**. Seedlings were grown for 7d on control plates and then transferred to control/Cd plates for 7d. Displayed are the root growth measurements 7d after transfer. Asterisks indicate statistical significance (p ≤ 0.05, Mann-Whitney U-test, data from three independent replications, n=15-30).

To further explore these findings, the Cd sensitivity of corresponding *A. thaliana* knock-out mutants was investigated. Phenotypic analysis of *elip1/2* and *hrg1* mutants revealed no significant difference in growth under Cd compared to the wild type (**Figure 4, B**). By contrast, Cd sensitivity was slightly but significantly higher for the mutants *xth20, myb12* and *hrg2.* Similarly, while *XTH26* was not specifically induced in RT by Cd, the corresponding mutant exhibited increased Cd susceptibility when compared to Col-0. The phenotype of an *xth12* mutant could not be investigated because no homozygous lines could be isolated from descendants of the segregating line SALK_008718. For *XTH13*, no appropriate line was available.

### Metabolomic analysis

To investigate metabolic changes upon Cd exposure in the two root parts, untargeted metabolomic analysis with LC-MS was conducted. The total content of 3052 detected features was remarkably different in RR and RT, as the RR samples displayed higher contents overall (**Figure 5, A**). Similar to the transcriptomic analysis, we found that the difference between RR and RT overshadowed the variance in metabolic profiles elicited by the treatment with Cd (**Figure 5, B**). Importantly, this global trend remains, when metabolite concentrations are normalized to the total sum of metabolites in each sample instead of a normalization based on the collected number of root pieces (**Figure 5, B, Supplementary Figure S3, A)**.

**Figure 5:**
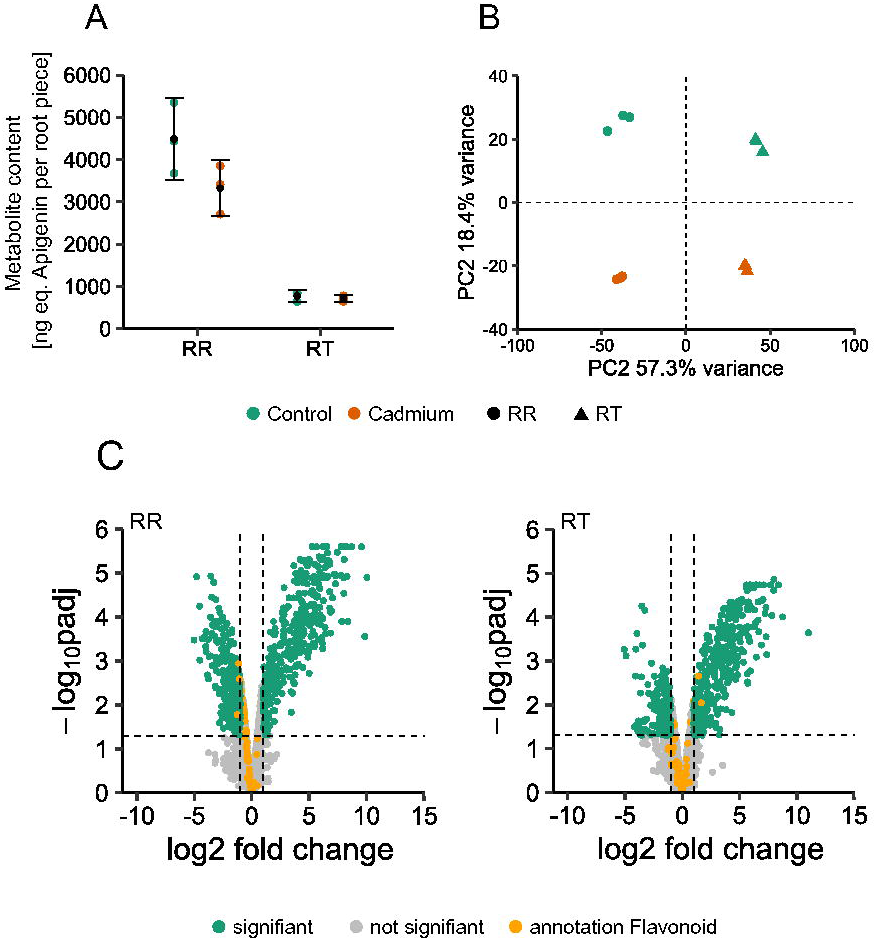
Metabolomic analysis of *A. thaliana* root tips (RT) and remaining roots (RR) 48h after exposure to 25µM Cd. **A.** Total content of all detected features in RR and RT samples. Data is presented as a relative quantification in ng equivalent to Apigenin (internal standard) per number of root pieces in each sample. Displayed are means +/- standard deviations. **B.** Principal component analysis showing the distribution of samples according to PC1 and PC2. The percentage of variance explained by each PC is indicated. Metabolite data were normalized to the total metabolite count in each sample **C.** Volcano plots showing the metabolic changes in RR and RT after Cd exposure. Significant differentially abundant metabolites (DAF) are indicated with green, non-significant DAF are labeled grey. Metabolites annotated as flavonoids are labeled orange. Statistical analysis was done with limma (p_adj_ ≤ 0.05). Only features displaying log2 fold changes greater or smaller than 1 were considered significant. Data are from three independent replications. Metabolite data were normalized to the total metabolite count in each sample.

The statistical analysis for differentially abundant features (DAF) revealed 164 features that were less abundant and 374 features that were more abundant after Cd treatment in RR (**Figure 5, C, left**). In RT, 175 DAF with lower abundance and 317 DAF with higher abundance due to Cd exposure were identified (**Figure 5, C, right**).

Phytochelatins (PCs) were detected with higher abundance in both RR and RT upon Cd exposure (**Supplementary Figure S3, D)**.

Since many GO terms related to irradiation were extracted (**Figure 2, C**), RT-specific upregulation of the flavonol-related transcription factor *MYB12* was observed and the *myb12* mutant displayed increased Cd sensitivity (**Figure 4, A, B**), differential abundance of features annotated as flavonoids was investigated. In RR, three features belonging to the flavonoid chemical family with lower concentration upon Cd exposure were identified. In RT, two flavonoid-related features were identified as significant DAF with higher concentration upon Cd exposure (**Figure 5, C**).

Consistent with the downregulation of *CYP82C4*, sideretin concentration decreased in RR after Cd exposure (**Supplementary Figure S3, C**).

### Cd accumulation and sensitivity of *hy5*

A literature search revealed that many of the genes displaying Cd-induced expression exclusively in RT, namely *XTH12, 13* and *20* (Bursch *et al*., 2020), *MYB12* (Stracke *et al*., 2010) and *ELIP1* and *2* (Burko *et al*., 2020) are regulated by the transcription factor HY5 (ELONGATED HYPOCOTYL 5). Furthermore, akin to *HRG1* and *HRG2,* a recent study has highlighted HY5’s involvement in ROS homeostasis within the root meristem (Gong *et al*., 2021; Li *et al*., 2024). Remarkably, under Cd exposure, a set of 297 high confidence HY5 target genes (Burko *et al*., 2020) demonstrated a collective upregulation clearly surpassing that of a background distribution generated by random sampling of *A. thaliana* genes (**Supplementary Figure S4**).

Phenotypic analysis of two different *hy5* mutants, named *hy5_1* and *hy5_215* (Oyama *et al*., 1997) in transfer experiments (7d control growth, 7d control/Cd25 growth) showed an enhanced sensitivity to Cd compared to the WT (**Figure 6, A**).

**Figure 6:**
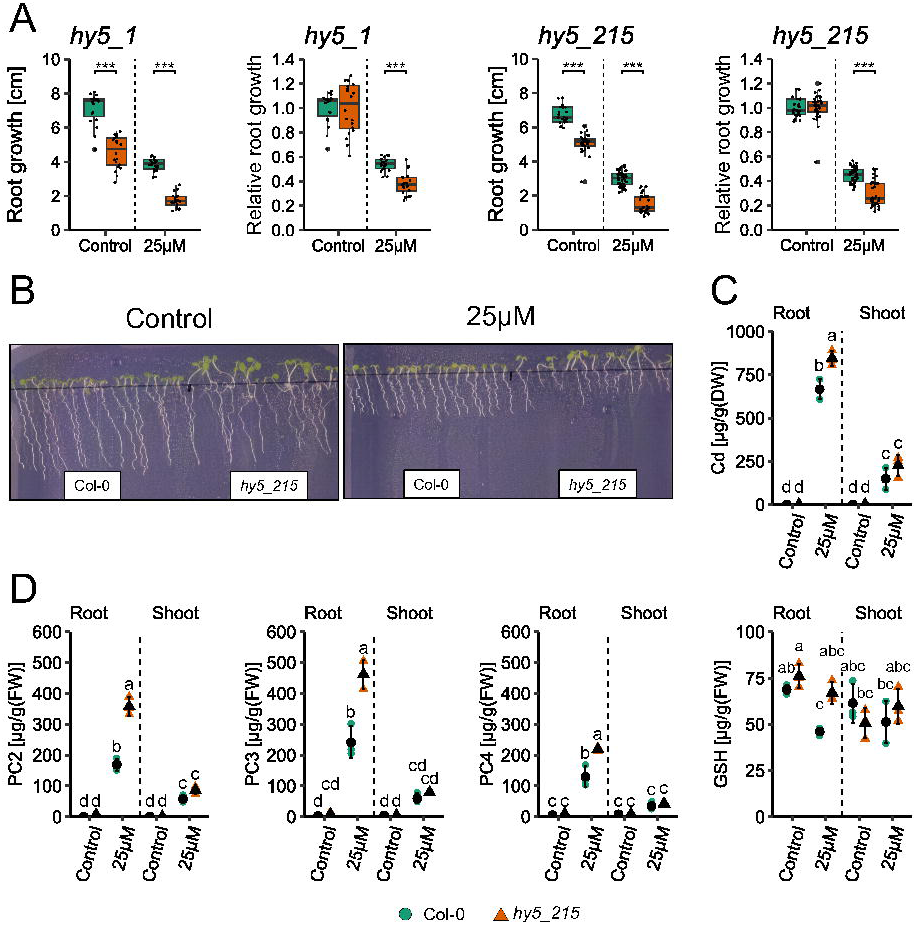
The phenotype of *hy5* under control conditions and 25µM Cd. **A.** Absolute and relative root growth of two *hy5* mutants under control and Cd conditions. Seedlings were grown for 7d on control plates and then transferred to control/Cd plates for 7d. Displayed are the root growth measurements 7d after transfer. Asterisks indicate statistical significance (p ≤ 0.05, Mann-Whitney U-test, data from three independent replications). **B.** Representative pictures of Col-0 and *hy5_215* seedlings grown for 9d directly on either control or Cd plates. **C.** Cd accumulation in Col-0 and *hy5_215*. Seedlings were grown for 7d on control plates, transferred to control/Cd plates and harvested after 3d. Displayed mean values +/- standard deviation of Cd contents as µg per gram dry weight (DW). Letters indicate statistical significance (ANOVA with Tukey HSD, p≤0.05, data from three independent replications) **D.** Phytochelatin (PC) and glutathione contents in Col-0 and *hy5_215*. Seedlings were grown for 9d on control plates and then exposed to Cd or control conditions for 3d. Displayed are mean values +/- standard deviations of contents in µg per gram fresh weight (FW). Letters indicate statistical significance (ANOVA with Tukey HSD, p≤0.05, data from three independent replications).

Because abundance of the HY5 protein is highly dependent on illumination (Zhang *et al*., 2019), we assessed the Cd phenotype of *hy5* mutants when roots were not lit. We found that both in a root-covered system (RCS) and in complete darkness, *hy5* mutants displayed stronger root growth reduction by Cd than the wild type, Col-0. Additionally, under these conditions, primary root growth in the absence of Cd was less affected in the mutants (**Supplementary Figures S5 and S6**).

Furthermore, when directly sown on control and Cd plates and grown for 9d, we observed very little difference in primary root length between Col-0 and *hy5_215* under control conditions. However, the increased Cd-sensitivity of the mutant was confirmed (**Figure 6, B**).

Metal concentrations in roots and shoots of the *hy5* mutants after Cd exposure were analyzed with ICP-OES. Cd concentrations in both roots and shoots of *hy5_215* were higher than in Col-0 (**Figure 6, C**). Similarly, elevated Cd concentrations in the *hy5_1* mutant were observed (**Supplementary Figure S1**). In agreement with the finding of increased Cd accumulation, both *hy5* mutants displayed increased PC contents (PC2, 3, 4) compared to wild type seedlings after Cd exposure (**Figure 6, D, Supplementary Figure S7**). For both Cd accumulation and PC contents, we found a higher increase in roots compared to shoots.

## Discussion

This study analyses the differential responses of *A. thaliana* root tips (RT) and remaining roots (RR) to Cd exposure. The impact of Cd on the RAM and EZ have previously been addressed (Yuan & Huang, 2016) and we confirmed a reduction in size of both root regions upon 48h of Cd stress (**Supplementary Figure S8**). The spatial separation of RT and RR enabled a specific analysis of the impact of metallic stress on the actively growing tissues in the RT. Additionally, our experimental setting also allowed us to compare the two root regions in control conditions, in particular their metal homeostasis.

### Reduced root growth is not accompanied by differential expression of cell cycle regulatory genes

One goal of this study was to analyze the specific impact of Cd on the expression of cell cycle regulatory genes in the RAM. Even though a reduction of primary root growth by Cd was observed after 24 and 48h (**Figure 1, A**), the transcriptomic data demonstrated that this was not accompanied by the differential expression of such genes (**Supplementary Figure S2**). Interestingly, a similar observation was made in RT of plants exposed to Zn excess (accompanying manuscript, Thiébaut *et al*.). This suggests that the mechanisms underlying Cd-induced root growth inhibition might not directly involve transcriptional alterations of cell cycle genes. Indeed, many actors of the cell cycle regulatory machinery, such as cyclin dependent kinases (CDKs) or cyclins, are known to be regulated at the post-translational level by mechanisms like phosphorylation and proteolysis (Inzé & Veylder, 2006). However, previous research on the cell cycle under Cd stress in RT described effects such as a reduction of the mitotic index (the ratio of cells in mitosis and the total cell number) or an altered expression of cell cycle-related genes (reviewed in Huybrechts *et al*., 2019). Noteworthy, the latter was reported exclusively in the context of prolonged exposure durations (*i.e.* > 5d). Thus, the transcriptional regulation of the cell cycle may occur only upon persistent Cd stress.

### Elemental concentrations in the two root parts are mirrored by transporter gene expression

After Cd treatment, RT showed lower Cd accumulation, with levels approximately half that of RR (**Figure 1, C**) and a lower impact on gene expression compared to RR (**Figure 2, A**).

Differences in the ionomic profiles between RR and RT were analyzed together with the expression of transporter genes. *IRT1*, the Fe(II) uptake transporter known for its role in Cd uptake in *A. thaliana* roots, was found to exhibit large differences in expression between the two root parts, which is in agreement with previous reports on the expression of *IRT1* in different root sections (Vert *et al*., 2002). Similarly to *IRT1,* the nitrate transporter *NRT2.1,* which also has a role in Cd uptake (Guan *et al*., 2021), was found to be less expressed in RT compared to RR. These differences in the expression of transporter genes may at least partially account for the reduced Cd concentrations observed in RT compared to RR. Importantly, lower concentrations of Zn in RT compared to RR, in parallel with lower expression of Zn uptake transporters, were also found upon exposure to Zn excess conditions (accompanying manuscript, Thiébaut *et al*.). These parallel observations for both ions support the possible presence of protective mechanisms that limit the entry of potentially toxic metals into the root apex.

To mitigate toxic effects, plants manage metals within the cytosol by chelation, exporting them to the apoplast or compartmentalizing them within suitable cellular locations such as the vacuole (Lin & Aarts, 2012). However, in the RAM, vacuoles are very small, and their size increases as cells transition into the EZ (Cui *et al*., 2019; Kaiser & Scheuring, 2020). Theoretically, the small size of vacuoles in the RAM and the associated lower storage capacity could also contribute to the observed differences in metal accumulation between RT and RR.

By contrast, higher expression of the transporter *COPT1*, which mediates copper acquisition (He *et al*., 2023), was found in RT compared to RR, which was consistent with the higher copper concentration measured in RT compared to RR (**Figure 1, B**).

One exception to the correlation of elemental concentrations with transporter gene expression in the two root parts concerns P, which was more abundant in RT (**Figure 1, B**). As a necessary component of nucleic acids, ATP and phospholipids, P is absorbed by plant roots in the form of orthophosphate (P_i_) mainly by the transporters PHT1;1 and PHT1;4 (Bucher, 2007; Yang *et al*., 2024), whose corresponding genes did not show significantly higher transcript levels in RT compared to RR (**Supplementary tables S5 and S6**). Nevertheless, the root tip was shown to be responsible for a substantial proportion of total P_i_ uptake in *A. thaliana*, with a high concentration of P_i_ transporters specifically at the root cap (Kanno *et al*., 2016).

### Cd triggers many established Fe deficiency responses but suppresses others

At the transcriptomic level, one prominent consequence of Cd exposure was the activation of transcriptomic changes similar to those observed under Fe deficiency (**Figure 2, C**). For instance, the repression of *FERRITIN* (Buckhout *et al*., 2009; Hantzis *et al*., 2018; von der Mark *et al*., 2020) and *VTL* (Gollhofer *et al*., 2014; Tabata *et al*., 2022) genes or the induction of *bHLH38/39* (Kim *et al*., 2019; Chen *et al*., 2021), *IRONMAN* (Grillet *et al*., 2018) and *FRO* (Riaz & Guerinot, 2021) genes were previously shown to occur under low Fe-availability and also under Cd exposure in this study (**Supplementary Tables S1-S4**). Similar to our ICP-OES data (**Supplementary Figure S1),** Fe concentrations were previously found to be decreased in *A. thaliana* roots upon Cd-exposure, possibly due to a competition for the shared uptake of the two metals by the low specificity Fe transporter IRT1 (Vert *et al*., 2002; Lešková *et al*., 2017). Especially because the activation of an Fe deficiency response under Cd is expected to counteract the systemic Fe deprivation caused by Cd, it is surprising that Cd exposure results in the downregulation of the *S8H* and *CYP82C4* genes, which are involved in the synthesis of catechol-type coumarins, along with a confirmed reduction of sideretin content (**Supplementary Figure S3, C**). Fe deficiency was shown to induce coumarins, Fe-mobilizing metabolites derived from the phenylpropanoid pathway. In particular, sideretin accumulates strongly under low Fe and specifically at acidic pH conditions such as the ones used in our experiments (Rajniak *et al*., 2018; Robe *et al*., 2021a; Vélez-Bermúdez & Schmidt, 2023). Regardless of these observations, the *cyp82c4* KO mutant used by Rajniak *et al.,* 2018 (Rajniak *et al*., 2018), lacking the ability to synthesize sideretin, showed wild type like growth under Cd-exposure (**Supplementary Figure S3, B**). Interestingly, evidence suggests an IRT1-independent pathway for the uptake of the catechol-type coumarin fraxetin complexed with Fe^3+^ (Robe *et al*., 2021b; Vélez-Bermúdez & Schmidt, 2022). While this phytosiderophore-like mechanism challenges the perception that IRT1 exclusively controls Fe entry into root cells, our data foster the speculation over a potential role of coumarins in Cd uptake. Notably, the finding that sideretin concentration and the expression of *CYP82C4* both increased in RR upon Zn excess (accompanying manuscript, Thiébaut *et al*.) also encourages the consideration of a mechanism for differentiation between essential and non-essential elements in coumarin-mediated metal uptake.

### HY5 affects Cd tolerance and accumulation

HY5, a transcription factor (TF) belonging to the basic leucine (bZIP) family, is known primarily for promoting photomorphogenesis, but also regulates various processes including circadian rhythm, ROS, hormone signaling, and nutrient uptake (Gangappa & Botto, 2016; Mankotia *et al*., 2024). Although HY5 was historically associated with light-regulated development, recent research has uncovered its role in root-specific processes, such as its translocation from shoots to roots and its autonomous expression in roots (Zhang et al., 2017, 2019; Burko et al., 2020a). However, the specific cofactors that associate with HY5 in roots and its target genes in these tissues remain unclear. Both transcriptional repression and induction of target genes have been associated with HY5 (Mankotia *et al*., 2024).

Our results reveal the involvement of HY5 in Cd tolerance, as two independent lines lacking functional HY5 exhibited greater root growth inhibition by Cd than wild type plants (**Figure 6, A, B**). This increased sensitivity was accompanied by an elevated Cd accumulation in roots (**Figure 6, C**), and by an increased formation of phytochelatins (PCs), compared to Col-0 (**Figure 6, D**). This indicates higher cytosolic Cd levels, since PC-synthases are activated upon elevated Cd levels inside cells (Blum *et al*., 2010).

Transcriptomic analysis identified several DEGs in RT under Cd stress that are regulated by HY5, including *MYB12*, *ELIP1/2*, *XTH12,13, 26*, and genes involved in ROS regulation, such as *HRG1/2* (Harari-Steinberg *et al*., 2001; Stracke *et al*., 2010; Bursch *et al*., 2020; Gong *et al*., 2021)(**Figure 4, A**).

The repression of X*TH12*, *13* and *26* in the absence of functional HY5 (Bursch *et al*., 2020) further underscores HY5’s role in regulating genes involved in cell wall modification under stress conditions.

Additionally, HY5’s involvement in ROS regulation is highlighted by the RT-specific upregulation of *HRG1* and *HRG2*, genes involved in the removal of H_2_O_2_ from the root apical meristem, which is crucial for maintaining meristem integrity and optimal root growth under stress (Gong *et al*., 2021).

Our metabolomic data also support the role of flavonoids in RT during Cd stress. In RT, three features annotated as flavonoids were significantly more abundant under Cd exposure. The increased Cd sensitivity of both *myb12* and *hy5* mutants, coupled with the regulation of *MYB12* by HY5 (Stracke *et al*., 2010), suggests that these transcription factors work together to synthesize specialized metabolites, potentially flavonoids, to maintain meristem integrity during Cd stress.

Moreover, the role of HY5 in sulfur assimilation is relevant to Cd accumulation, as it directly regulates *APR1* and *APR2*, key genes in sulfur metabolism (Lee et al., 2011; Mankotia et al., 2024). The *hy5* mutants exhibited higher levels of GSH and PCs, which may contribute to increased Cd accumulation (**Figure 6, D**; **Supplementary Figure S7**).

Changes in root architecture, such as increased root hair length and lateral root density in *hy5* mutants, may also enhance Cd absorption. However, increased sensitivity towards Cd was observed across various light conditions **(Supplementary Figures S5 and S6**) and changes in illumination drastically impact root morphology (Miotto *et al*., 2021; Cabrera *et al*., 2022), pointing to altered uptake mechanisms as a more likely cause.

HY5 also plays a role in Fe uptake, and our findings suggest that disruption of Fe homeostasis in *hy5* mutants under Cd stress contributes to increased Cd accumulation. While Fe concentrations decreased in Col-0 roots under Cd stress, they remained stable in *hy5* mutants (**Supplementary Figure S1)**, indicating disrupted Fe uptake regulation.

This study uncovers a novel role for HY5 in limiting Cd accumulation, adding to its documented functions in nutrient signaling and uptake. Together with other studies, these findings underscore the importance of central regulators like HY5 in plant responses to environmental stress.

## Acknowledgements

We thank L. Karim and Dr. W. Coppieters (ULiège, Belgium) for performing the RNA sequencing, Dr. M. Corso (Institut Jean-Pierre Bourgin, France) for support in metabolomics data analyses and Sarah Plößner for assisting in the HPLC measurements. We thank Prof. L. Veylder (UGent, Belgium) for helpful discussion at the start of the project. We thank Bert De Rybel (Ghent University, Belgium) and Andras Viczian (University of Szeged, Hungary) for providing the seeds of *myb12-1-f* and *hy5_215*, respectively. Funding was provided by the “Fonds de la Recherche Scientifique-FNRS” (PDR-T0120.18, PDR-T.0104.22 to M.H. and N.V.), the COST ACTION 19116 PLANTMETALS, and the University of Bayreuth (to SC). The IJPB benefits from the support of Saclay Plant Sciences-SPS (ANR-17-EUR-0007). This work has benefited from the support of IJPB’s Plant Observatory platform PO-Chem.

## Competing interests

The authors declare no competing interests.

## Author contributions

NV, SC, LR, NT and MH designed the research. LR, NT, AR and SB performed experiments. LR, NT, SB, NV, MH, SC and MS analyzed the data. LR, NT and MS made the figures. LR, SC, NV, MH, NT and MS wrote the manuscript. All authors read and approved the manuscript.

## Data availability

The RNA-Seq reads have been deposited in the National Center for Biotechnology Information (NCBI) Sequence Read Archive (SRA) Database with BioProject (submission in progress). The metabolomic data and metadata have been deposited at the MassiVE data repository portal (submission in progress). The other data that support the findings of this study are available from the corresponding authors upon reasonable request.

## Supporting Information

**Figure S1.** ICP-OES measurement of Cd (left) and Fe (right) in Col-0 and *hy5_1* seedlings upon exposure to 25µM Cd.

**Figure S2.** Effect of 25µM Cd on expression of cell cycle related genes from Vandepoele et al. 2002 in root tips (RT) after 24h and 48h.

**Figure S3.** The regulation of specialized metabolism in response to 25µM Cd.

**Figure S4.** Regulation of 297 direct HY5 target genes from Burko et al 2020 in comparison to random distributions upon Cd exposure.

**Figure S5.** Primary root growth of *hy5_1* and *hy5_215* under exposure to 25µM Cd in a root covered system.

**Figure S6.** Primary root growth of *hy5_1* under exposure to 25µM Cd in darkness.

**Figure S7.** Phytochelatin (PC) and glutathione contents in Col-0 and *hy5_1* upon exposure to 25µM Cd.

**Figure S8.** Effects of 25µM Cd on the root apical meristem (RAM) and elongation zone (EZ).

**Figure S9.** Primary root growth of Col-0 *and hy5_215* under different Fe concentrations.

**Table S1.** DEG in RR after 24h of Cd exposure.

**Table S2.** DEG in RR after 48h of Cd exposure.

**Table S3.** DEG in RT after 24h of Cd exposure.

**Table S4.** DEG in RT after 48h of Cd exposure.

**Table S5.** Genes less expressed in RR compared to RT.

**Table S6.** Genes more expressed in RR compared to RT.

## Notes

### Competing Interest Statement

The authors have declared no competing interest.

## Literature

Bahmani R, Kim D, Modareszadeh M, Hwang S. 2022. Cadmium enhances root hair elongation through reactive oxygen species in Arabidopsis. Environmental and Experimental Botany 196: 104813.

Blum R, Meyer KC, Wünschmann J, Lendzian KJ, Grill E. 2010. Cytosolic Action of Phytochelatin Synthase. Plant Physiology 153: 159–169.

Bolger AM, Lohse M, Usadel B. 2014. Trimmomatic: a flexible trimmer for Illumina sequence data. *Bioinformatics (Oxford*, England) 30: 2114–2120.

Boutet S, Barreda L, Perreau F, Totozafy J-C, Mauve C, Gakière B, Delannoy E, Martin-Magniette M-L, Monti A, Lepiniec L, et al. 2022. Untargeted metabolomic analyses reveal the diversity and plasticity of the specialized metabolome in seeds of different Camelina sativa genotypes. The Plant Journal: For Cell and Molecular Biology 110: 147–165.

Bruno L, Pacenza M, Forgione I, Lamerton LR, Greco M, Chiappetta A, Bitonti MB. 2017. In Arabidopsis thaliana Cadmium Impact on the Growth of Primary Root by Altering SCR Expression and Auxin-Cytokinin Cross-Talk. Frontiers in Plant Science 8.

Bucher M. 2007. Functional biology of plant phosphate uptake at root and mycorrhiza interfaces. New Phytologist 173: 11–26.

Buckhout TJ, Yang TJ, Schmidt W. 2009. Early iron-deficiency-induced transcriptional changes in Arabidopsis roots as revealed by microarray analyses. BMC Genomics 10: 147.

Burko Y, Seluzicki A, Zander M, Pedmale UV, Ecker JR, Chory J. 2020. Chimeric Activators and Repressors Define HY5 Activity and Reveal a Light-Regulated Feedback Mechanism[OPEN]. The Plant Cell 32: 967–983.

Bursch K, Toledo-Ortiz G, Pireyre M, Lohr M, Braatz C, Johansson H. 2020. Identification of BBX proteins as rate-limiting cofactors of HY5. Nature Plants 6: 921–928.

Cabrera J, Conesa CM, del Pozo JC. 2022. May the dark be with roots: a perspective on how root illumination may bias in vitro research on plant–environment interactions. New Phytologist 233: 1988–1997.

Chen W, Zhao L, Liu L, Li X, Li Y, Liang G, Wang H, Yu D. 2021. Iron deficiency-induced transcription factors bHLH38/100/101 negatively modulate flowering time in *Arabidopsis thaliana*. Plant Science 308: 110929.

Cheng C-Y, Krishnakumar V, Chan AP, Thibaud-Nissen F, Schobel S, Town CD. 2017. Araport11: a complete reannotation of the Arabidopsis thaliana reference genome. The Plant Journal 89: 789–804.

Clemens S, Aarts MGM, Thomine S, Verbruggen N. 2013. Plant science: the key to preventing slow cadmium poisoning. Trends in Plant Science 18: 92–99.

Connolly EL, Fett JP, Guerinot ML. 2002. Expression of the IRT1 Metal Transporter Is Controlled by Metals at the Levels of Transcript and Protein Accumulation. The Plant Cell 14: 1347–1357.

Cui Y, Cao W, He Y, Zhao Q, Wakazaki M, Zhuang X, Gao J, Zeng Y, Gao C, Ding Y, et al. 2019. A whole-cell electron tomography model of vacuole biogenesis in Arabidopsis root cells. Nature Plants 5: 95–105.

Cui W, Wang H, Song J, Cao X, Rogers HJ, Francis D, Jia C, Sun L, Hou M, Yang Y, et al. 2017. Cell cycle arrest mediated by Cd-induced DNA damage in Arabidopsis root tips. Ecotoxicology and Environmental Safety 145: 569–574.

Dello Ioio R, Linhares FS, Scacchi E, Casamitjana-Martinez E, Heidstra R, Costantino P, Sabatini S. 2007. Cytokinins Determine *Arabidopsis* Root-Meristem Size by Controlling Cell Differentiation. Current Biology 17: 678–682.

Fan SK, Fang XZ, Guan MY, Ye YQ, Lin XY, Du ST, Jin CW. 2014. Exogenous abscisic acid application decreases cadmium accumulation in Arabidopsis plants, which is associated with the inhibition of IRT1-mediated cadmium uptake. Frontiers in Plant Science 5.

Gangappa SN, Botto JF. 2016. The Multifaceted Roles of HY5 in Plant Growth and Development. Molecular Plant 9: 1353–1365.

Gollhofer J, Timofeev R, Lan P, Schmidt W, Buckhout TJ. 2014. Vacuolar-Iron-Transporter1-Like Proteins Mediate Iron Homeostasis in Arabidopsis. PLOS ONE 9: e110468.

Gong F, Yao Z, Liu Y, Sun M, Peng X. 2021. H2O2 response gene 1/2 are novel sensors or responders of H2O2 and involve in maintaining embryonic root meristem activity in *Arabidopsis thaliana*. Plant Science 310: 110981.

Grillet L, Lan P, Li W, Mokkapati G, Schmidt W. 2018. IRON MAN is a ubiquitous family of peptides that control iron transport in plants. Nature Plants 4: 953–963.

Guan M, Chen M, Cao Z. 2021. NRT2.1, a major contributor to cadmium uptake controlled by high-affinity nitrate transporters. Ecotoxicology and Environmental Safety 218: 112269.

Haider FU, Liqun C, Coulter JA, Cheema SA, Wu J, Zhang R, Wenjun M, Farooq M. 2021. Cadmium toxicity in plants: Impacts and remediation strategies. Ecotoxicology and Environmental Safety 211: 111887.

Hantzis LJ, Kroh GE, Jahn CE, Cantrell M, Peers G, Pilon M, Ravet K. 2018. A Program for Iron Economy during Deficiency Targets Specific Fe Proteins. Plant Physiology 176: 596– 610.

Harari-Steinberg O, Ohad I, Chamovitz DA. 2001. Dissection of the Light Signal Transduction Pathways Regulating the Two Early Light-Induced Protein Genes in Arabidopsis. Plant Physiology 127: 986–997.

He L, Ma H, Song W, Zhou Z, Ma C, Zhang H. 2023. Arabidopsis COPT1 copper transporter uses a single histidine to regulate transport activity and protein stability. International Journal of Biological Macromolecules 241: 124404.

Howden R, Goldsbrough PB, Andersen CR, Cobbett CS. 1995. Cadmium-Sensitive, cad1 Mutants of Arabidopsis thaliana Are Phytochelatin Deficient. Plant Physiology 107: 1059–1066.

Huybrechts M, Cuypers A, Deckers J, Iven V, Vandionant S, Jozefczak M, Hendrix S. 2019. Cadmium and Plant Development: An Agony from Seed to Seed. International Journal of Molecular Sciences 20: 3971.

Inzé D, Veylder LD. 2006. Cell Cycle Regulation in Plant Development1. Annual Review of Genetics 40: 77–105.

Ishimaru Y, Takahashi R, Bashir K, Shimo H, Senoura T, Sugimoto K, Ono K, Yano M, Ishikawa S, Arao T, et al. 2012. Characterizing the role of rice NRAMP5 in Manganese, Iron and Cadmium Transport. Scientific Reports 2: 286.

Ismael MA, Elyamine AM, Moussa MG, Cai M, Zhao X, Hu C. 2019. Cadmium in plants: uptake, toxicity, and its interactions with selenium fertilizers. Metallomics 11: 255–277.

Jia Z, Giehl RFH, Hartmann A, Estevez JM, Bennett MJ, Wirén N von. 2023. A spatially concerted epidermal auxin signaling framework steers the root hair foraging response under low nitrogen. Current Biology 33: 3926–3941.e5.

Kaiser S, Scheuring D. 2020. To Lead or to Follow: Contribution of the Plant Vacuole to Cell Growth. Frontiers in Plant Science 11.

Kanno S, Arrighi J-F, Chiarenza S, Bayle V, Berthomé R, Péret B, Javot H, Delannoy E, Marin E, Nakanishi TM, et al. 2016. A novel role for the root cap in phosphate uptake and homeostasis (D Weigel, Ed.). eLife 5: e14577.

Kim SA, LaCroix IS, Gerber SA, Guerinot ML. 2019. The iron deficiency response in Arabidopsis thaliana requires the phosphorylated transcription factor URI. Proceedings of the National Academy of Sciences 116: 24933–24942.

Kumar S, Kumar S, Mohapatra T. 2021. Interaction Between Macro- and Micro-Nutrients in Plants. Frontiers in Plant Science 12: 665583.

Lešková A, Giehl RFH, Hartmann A, Farga¡ová A, von Wirén N. 2017. Heavy Metals Induce Iron Deficiency Responses at Different Hierarchic and Regulatory Levels. Plant Physiology 174: 1648–1668.

Leonardo B, Emanuela T, Letizia MM, Antonella M, Marco M, Fabrizio A, Beatrice BM, Adriana C. 2021. Cadmium affects cell niches maintenance in *Arabidopsis thaliana* post-embryonic shoot and root apical meristem by altering the expression of WUS/WOX homolog genes and cytokinin accumulation. Plant Physiology and Biochemistry 167: 785–794.

Li J, Zeng J, Tian Z, Zhao Z. 2024. Root-specific photoreception directs early root development by HY5-regulated ROS balance. Proceedings of the National Academy of Sciences 121: e2313092121.

Lin Y-F, Aarts MGM. 2012. The molecular mechanism of zinc and cadmium stress response in plants. Cellular and Molecular Life Sciences 69: 3187–3206.

Love MI, Huber W, Anders S. 2014. Moderated estimation of fold change and dispersion for RNA-seq data with DESeq2. Genome Biology 15: 550.

Lux A, Martinka M, Vaculík M, White PJ. 2011. Root responses to cadmium in the rhizosphere: a review. Journal of Experimental Botany 62: 21–37.

Mankotia S, Jakhar P, Satbhai SB. 2024. HY5: a key regulator for light-mediated nutrient uptake and utilization by plants. New Phytologist 241: 1929–1935.

von der Mark C, Ivanov R, Eutebach M, Maurino VG, Bauer P, Brumbarova T. 2020. Reactive oxygen species coordinate the transcriptional responses to iron availability in Arabidopsis. Journal of Experimental Botany 72: 2181–2195.

Mehrtens F, Kranz H, Bednarek P, Weisshaar B. 2005. The Arabidopsis Transcription Factor MYB12 Is a Flavonol-Specific Regulator of Phenylpropanoid Biosynthesis. Plant Physiology 138: 1083–1096.

Miotto YE, da Costa CT, Offringa R, Kleine-Vehn J, Maraschin F dos S. 2021. Effects of Light Intensity on Root Development in a D-Root Growth System. Frontiers in Plant Science 12: 778382.

Oyama T, Shimura Y, Okada K. 1997. The Arabidopsis HY5 gene encodes a bZIP protein that regulates stimulus-induced development of root andc:hypocotyl. Genes & Development 11: 2983–2995.

Perilli S, Di Mambro R, Sabatini S. 2012. Growth and development of the root apical meristem. Current Opinion in Plant Biology 15: 17–23.

Perilli S, Moubayidin L, Sabatini S. 2010. The molecular basis of cytokinin function. Current Opinion in Plant Biology 13: 21–26.

Pischke E, Barozzi F, Colina Blanco AE, Kerl CF, Planer-Friedrich B, Clemens S. 2022. Dimethylmonothioarsenate Is Highly Toxic for Plants and Readily Translocated to Shoots. Environmental Science & Technology 56: 10072–10083.

Rahim HU, Akbar WA, Alatalo JM. 2022. A Comprehensive Literature Review on Cadmium (Cd) Status in the Soil Environment and Its Immobilization by Biochar-Based Materials. Agronomy 12: 877.

Rajniak J, Giehl RFH, Chang E, Murgia I, von Wirén N, Sattely ES. 2018. Biosynthesis of redox-active metabolites in response to iron deficiency in plants. Nature chemical biology 14: 442–450.

Ren M, Li Y, Zhu J, Zhao K, Wu Z, Mao C. 2023. Phenotypes and Molecular Mechanisms Underlying the Root Response to Phosphate Deprivation in Plants. International Journal of Molecular Sciences 24: 5107.

Riaz N, Guerinot ML. 2021. All together now: regulation of the iron deficiency response. Journal of Experimental Botany 72: 2045–2055.

Robe K, Conejero G, Gao F, Lefebvre-Legendre L, Sylvestre-Gonon E, Rofidal V, Hem S, Rouhier N, Barberon M, Hecker A, et al. 2021a. Coumarin accumulation and trafficking in Arabidopsis thaliana: a complex and dynamic process. New Phytologist 229: 2062–2079.

Robe K, Stassen M, Chamieh J, Gonzalez P, Hem S, Santoni V, Dubos C, Izquierdo E. 2021b. Uptake of Fe-fraxetin complexes, an IRT1 independent strategy for iron acquisition in Arabidopsis thaliana. : 2021.08.03.454955.

Růžička K, Šimášková M, Duclercq J, Petrášek J, Zažímalová E, Simon S, Friml J, Van Montagu MCE, Benková E. 2009. Cytokinin regulates root meristem activity via modulation of the polar auxin transport. Proceedings of the National Academy of Sciences 106: 4284– 4289.

Spielmann J, Ahmadi H, Scheepers M, Weber M, Nitsche S, Carnol M, Bosman B, Kroymann J, Motte P, Clemens S, et al. 2020. The two copies of the zinc and cadmium ZIP6 transporter of Arabidopsis halleri have distinct effects on cadmium tolerance. Plant, Cell & Environment 43: 2143–2157.

Stracke R, Favory J-J, Gruber H, Barllniewoehner L, Barlls S, Binkert M, Funk M, Weisshaar B, Ulm R. 2010. The Arabidopsis bZIP transcription factor HY5 regulates expression of the PFG1/MYB12 gene in response to light and ultraviolet-B radiation. Plant, Cell & Environment 33: 88–103.

Tabata R, Kamiya T, Imoto S, Tamura H, Ikuta K, Tabata M, Hirayama T, Tsukagoshi H, Tanoi K, Suzuki T, et al. 2022. Systemic Regulation of Iron Acquisition by Arabidopsis in Environments with Heterogeneous Iron Distributions. Plant and Cell Physiology 63: 842–854.

Tao L, Zhu H, Luo X, Li J, Ru Y, Lv J, Pan W, Li Y, Li X, Chen Y, et al. 2024. Manganese toxicity elicits the degradation of auxin transport carriers to restrain arabidopsis root growth. Environmental and Experimental Botany 225: 105863.

Thiébaut N, Richtmann L, Sarthou M, Persson DP, Ranjan A, Schloesser M, Boutet S, Rezende L, Assunção A, Clemens S, Verbruggen N, Hanikenne M. Specific redox and iron homeostasis responses in the root tip of Arabidopsis upon zinc excess. Submitted as companion paper to New Phytologist.

Tsukagoshi H, Busch W, Benfey PN. 2010. Transcriptional Regulation of ROS Controls Transition from Proliferation to Differentiation in the Root. Cell 143: 606–616.

Vandepoele K, Raes J, De Veylder L, Rouzé P, Rombauts S, Inzé D. 2002. Genome-Wide Analysis of Core Cell Cycle Genes in Arabidopsis. The Plant Cell 14: 903–916.

Vélez-Bermúdez IC, Schmidt W. 2022. How Plants Recalibrate Cellular Iron Homeostasis. Plant and Cell Physiology 63: 154–162.

Vélez-Bermúdez IC, Schmidt W. 2023. Plant strategies to mine iron from alkaline substrates. Plant and Soil 483: 1–25.

Vert G, Grotz N, Dédaldéchamp F, Gaymard F, Guerinot ML, Briat J-F, Curie C. 2002. IRT1, an Arabidopsis Transporter Essential for Iron Uptake from the Soil and for Plant Growth. The Plant Cell 14: 1223–1233.

Wang R, Fei Y, Pan Y, Zhou P, Adegoke JO, Shen R, Lan P. 2023. IMA peptides function in iron homeostasis and cadmium resistance. Plant Science: An International Journal of Experimental Plant Biology 336: 111868.

Wang H-Q, Xuan W, Huang X-Y, Mao C, Zhao F-J. 2021. Cadmium Inhibits Lateral Root Emergence in Rice by Disrupting OsPIN-Mediated Auxin Distribution and the Protective Effect of OsHMA3. Plant and Cell Physiology 62: 166–177.

Wells DM, Wilson MH, Bennett MJ. 2010. Feeling UPBEAT about Growth: Linking ROS Gradients and Cell Proliferation. Developmental Cell 19: 644–646.

Wu H, Chen C, Du J, Liu H, Cui Y, Zhang Y, He Y, Wang Y, Chu C, Feng Z, et al. 2012. Co-Overexpression FIT with AtbHLH38 or AtbHLH39 in Arabidopsis-Enhanced Cadmium Tolerance via Increased Cadmium Sequestration in Roots and Improved Iron Homeostasis of Shoots. Plant Physiology 158: 790–800.

Yang S-Y, Lin W-Y, Hsiao Y-M, Chiou T-J. 2024. Milestones in understanding transport, sensing, and signaling of the plant nutrient phosphorus. The Plant Cell 36: 1504–1523.

Yuan H-M, Huang X. 2016. Inhibition of root meristem growth by cadmium involves nitric oxide-mediated repression of auxin accumulation and signalling in Arabidopsis. Plant, Cell & Environment 39: 120–135.

Zhai Z, Gayomba SR, Jung H, Vimalakumari NK, Piñeros M, Craft E, Rutzke MA, Danku J, Lahner B, Punshon T, et al. 2014. OPT3 Is a Phloem-Specific Iron Transporter That Is Essential for Systemic Iron Signaling and Redistribution of Iron and Cadmium in Arabidopsis. The Plant Cell 26: 2249–2264.

Zhang Y, Wang C, Xu H, Shi X, Zhen W, Hu Z, Huang J, Zheng Y, Huang P, Zhang K-X, et al. 2019. HY5 Contributes to Light-Regulated Root System Architecture Under a Root-Covered Culture System. Frontiers in Plant Science 10.

Zhou M, Zhang LL, Ye JY, Zhu QY, Du WX, Zhu YX, Liu XX, Lin XY, Jin CW. 2021. Knockout of *FER* decreases cadmium concentration in roots of *Arabidopsis thaliana* by inhibiting the pathway related to iron uptake. Science of The Total Environment 798: 149285.

